# Temporal and Spatial Analysis of Event-Related Potentials in Response to Color Saliency Differences Among Various Color Vision Types

**DOI:** 10.1101/2023.09.12.557493

**Authors:** Naoko Takahashi, Masataka Sawayama, Xu Chen, Yuki Motomura, Hiroshige Takeichi, Satoru Miyauchi, Chihiro Hiramatsu

## Abstract

Individuals with minority color vision phenotypes have been reported to exhibit enhanced color discrimination and color recognition, which deviate from predictions based on their receptoral sensitivities. However, the specific mechanisms underlying this enhanced sensitivity remain unclear. In this study, we examined the commonality and diversity of neural activity between typical and anomalous trichromats in response to differences in color saliency. Electroencephalography was recorded during an oddball task, in which participants discriminated each of two target stimuli, blue-green and red, from a green standard stimulus. The chromaticity of the stimulus was identical across participants, whereas the relationship of saliency between the target stimuli was expected to be reversed between color vision types. The spatiotemporal dynamics of event-related potentials (ERPs) were analyzed using cluster-based permutation tests. Typical trichromats demonstrated faster behavioral and neural responses to the more salient red target stimulus, with pronounced neural activity spreading from the occipital to the parietal regions in the comparison between the target stimuli. Anomalous trichromats also exhibited similar temporal patterns toward the more salient target stimulus when each target stimulus was compared with the green standard stimulus, indicating comparable processing toward saliency across color vision types. Although a similarity was observed, neither behavioral nor neural responses in anomalous trichromats reflected saliency contrast differences. In addition, a comparative analysis of ERPs between color vision types did not reveal any distinct differences in either target stimulus. Given the large variations in color sensitivity in individuals with anomalous trichromacy, further investigation is required to understand the detailed neural processing in individuals with various color vision types.

## INTRODUCTION

Color perception is a subjective experience unique to each individual. Colors serve diverse functions, including aiding in object detection and recognition for prompt environmental understanding (Conway et al., 2023) and enriching the qualitative aspects of perception by adding aesthetic values and/or conveying symbolic meanings (Elliot and Maier, 2007; Muratbekova and Shamoi, 2024). Despite their internal nature, colors are communicated to others through color names in everyday discourse (Berlin and Kay, 1968; Kay and Regier, 2003). However, sharing subjective experiences can be more abstract, particularly when individuals with diverse sensory characteristics are involved (Hiramatsu et al., 2023). Therefore, characterizing the neural mechanisms underlying diverse color experiences across various types of color vision remains an important challenge in the field of visual neuroscience.

Variations in color vision are a common genetic trait in human population, with a small percentage of individuals (predominantly males due to the X-chromosome linked inheritance of color vision) exhibiting minor sensory characteristics of S, M, and L cone photoreceptor cells that differs from the prevalent type of trichromacy, hereafter referred to as ‘typical trichromacy’ (Birch, 2012; Neitz and Neitz, 2011; Deeb, 2005; Asenjo et al., 1994; Nathans et al., 1986). The majority of these variations can be attributed to altered photosensitivity, particularly in the L and M cone cells. Among minority color vision phenotypes, approximately 6 % of Caucasian males (Birch, 2012) and 3 % of Asian males (Okajima, 2011) have anomalous trichromacy, owing to a photosensitivity shift in a single class of cone cells, while approximately 2 % have dichromatic color vision based on two classes of cone cells.

Variations in the L and M cone sensitivities result in confined red-green color discrimination. The responses of the L and M cone cells are compared early in visual processing, and the relative outputs of these cone contrasts form the basis of the cardinal red-green color axis (Werner and Wooten, 1979). Generally, sensitivity shifts diminish the contrast gains from the L/M comparison, thereby reducing the perceived chromatic differences (Pokorny and Smith, 1977). In anomalous trichromacy, the spectral sensitivity separation of the L and M cones ranges between 1 and 12 nm (Merbs and Nathans, 1992; Asenjo et al., 1994; Neitz and Neitz, 2011), whereas the separation in typical trichromacy is approximately 25 nm (Dartnall et al., 1983; Merbs and Nathans, 1992; Stockman and Sharpe, 2000). In dichromacy, the absence of the L/M contrast results in the absence of the red-green color axis, and color vision is theoretically based on the blue-yellow axis derived from S cone responses, in contrast to the L or M cone cells. However, the actual color-discrimination capacities of individuals with anomalous trichromacy and dichromacy do not always align with the anticipated color perception based on their specific cone sensitivities (Bosten, 2019). A higher red-green sensitivity than that expected from cone sensitivity has been reported in a wide range of studies, including psychophysical color-matching experiments and color-naming tasks with cognitive components (Scheibner and Boynton, 1968; Smith and Pokorny, 1977; Nagy and Boynton, 1979; Montag, 1994; Neitz et al., 1999). For example, in a color-naming experiment, dichromats exhibited color category assessment capabilities similar to typical trichromacy when color stimuli were presented with sufficient size and time for identification (Montag, 1994).

Several possible mechanisms have been suggested to explain the extended red-green sensitivity, such as: rod contribution providing alternative signals (Smith and Pokorny, 1977; Nagy and Boynton, 1979; Montag and Boynton, 1987), enhancement of S-cone sensitivity in long-wavelength regions serving as alternative signals (Scheibner and Boynton, 1968), and variation in optical density (Neitz et al., 1999; Thomas et al., 2011). A more recent psychophysical study suggested post-receptoral enhancement (Boehm et al., 2021), while one using fMRI reported enhanced neural activity in the early visual cortex (V2) during anomalous trichromacy, even during simple fixation tasks (Tregillus et al., 2021). Overall, current evidence suggests that multiple mechanisms contribute to the extended sensitivity observed in anomalous trichromacy and dichromacy. However, despite past findings, the complete picture remains unclear. Studies linking perceptual and cognitive diversity to neural diversity are limited.

Therefore, we aimed to characterize the patterns of spatiotemporal neural activity in different color vision types during a sequence of neural activities involved in perceptual cognitive processing. Specifically, we investigated the neural activity measured by electroencephalography (EEG) during an attention-demanding task, as attention is an indispensable cognitive mechanism used in everyday visual tasks where chromatic differences often serve as cues for distinguishing objects from their background. Using chromatically identical stimuli for all participants allowed us to directly compare the effects of differences in chromatic sensitivity across various color vision types.

## 2 MATERIAL AND METHOD

### 2.1 Participants

Nineteen male participants with normal or corrected-to-normal visual acuity participated in this study, the majority of whom were undergraduate and graduate students at Kyushu University School of Design (mean age ± standard deviation : 23.32 ± 2.68 years). Participation was restricted to males, as approximately 10 % of females carry variant red and green visual pigment genes, which can modify cone sensitivity, with uncertain effects on color perception (Jordan et al., 2010).

To assess each participant’s color vision type, four color vision tests were conducted, including the Ishihara pseudoisochromatic plate test, Colour Assessment and Diagnosis (CAD) test (Barbur et al., 2021), HMC-anomaloscope (Oculus), and Farnsworth–Munsell 100 hue test. Based on the combined results of these tests, five participants were identified as deuteranomalous trichromats (anomalous trichromacy with altered sensitivity in the M cone), and one as a deuteranope (dichromacy without the M cone), while the remaining were typical trichromats.

Each participant was provided a financial compensation for their participation. The experimental procedure complied with the Declaration of Helsinki, and was approved by the Ethics Committee of the Graduate School of Design at Kyushu University (Approval No. 316). Written informed consent was obtained from each participant prior to the experiment.

To consider the perceptual variability in the color vision of each individual, red-green color sensitivity was measured using the CAD test. Table **??** shows the measured red-green thresholds of the deuteranomalous and deuteranope participants. These thresholds represent the amount of saturation required to perceive the color from the neutral gray point along the evaluation color axis, where a threshold of 1 represents the average discrimination threshold of typical trichromats (Barbur and Rodriguez-Carmona, 2017; Barbur et al., 2021).

The average threshold for typical trichromats in this study was 1.1 *±* 0.2, whereas that for anomalous trichromats was 13.1 ± 7.9.

### 2.2 Stimuli

A set of color stimuli was used to convey different chromatic contrasts when the two color stimuli were paired with a common standard stimulus. The chromaticities of the stimuli were selected based on the CIE 1976 u’, v’ uniform chromaticity scale diagram, allowing for the estimation of perceptual chromatic distances among the stimuli (Pointer, 1981). Three colors; blue-green, orange-tinted red (hereafter referred to as red), and green, were selected to be equidistant from the neutral gray of D65 in the u’, v’ diagram (Figure 1). The specific coordinates of the stimuli were (0.1679, 0.4670) for blue-green, (0.2278, 0.4696) for red, and (0.1817, 0.4936) for green, each situated at an equal distance (0.03) from D65 (0.1978, 0.4683), to ensure that each stimulus had equal saturation relative to D65. In the attention task, the target (deviant) stimuli were blue-green or red, in contrast to the standard green stimulus.

**Figure 1.**
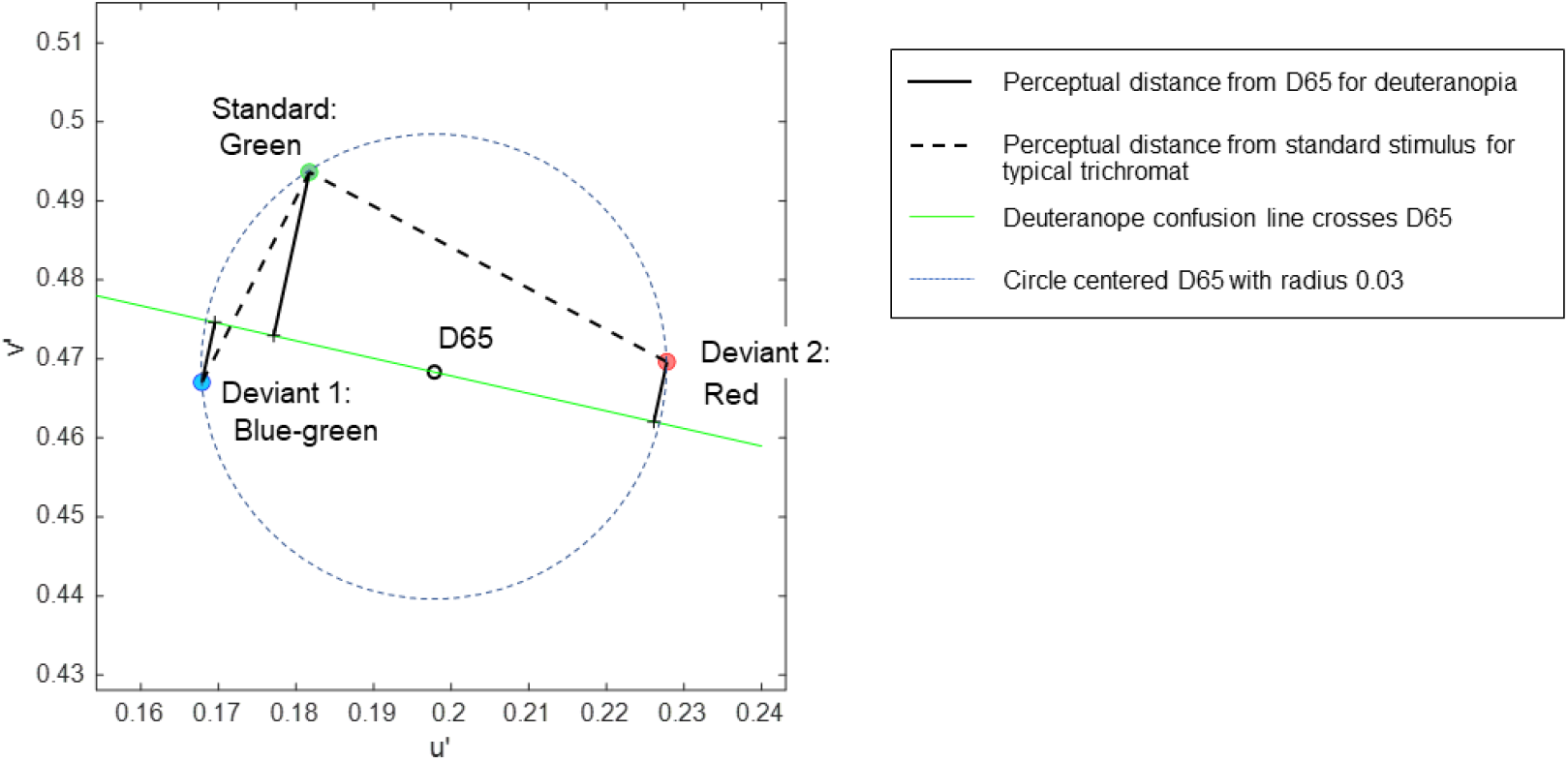
The coordinates of stimuli in the CIE 1976 u’, v’ uniform chromaticity scale diagram, in which perceptual distance is represented in terms of the Euclidean distance for typical trichromats. The expected chromatic contrasts between the standard and deviant stimuli are represented as the length of the lines for typical trichromacy (black dotted lines) and deuteranopia (black solid lines).

In the uniform color space, perceptual differences in colors are represented as Euclidean distances for typical trichromacy; the greater the distance between two points, the larger the difference in perception. The Euclidean distances of blue-green and red from green in the space were 0.03 and 0.052, respectively. For typical trichromacy, the red pair was expected to be more salient than the blue-green pair because the Euclidean distance from standard green was greater for red than for blue-green. This relationship can also be explained by categorical color differences: blue-green and green are expected to be categorically more similar than red and green in typical trichromacy. Thus, visual saliency was expected to be greater for red than blue-green when contrasted with green.

The color space of minority color vision phenotypes is still debated (Broackes, 2010). However, theoretical estimates of relative chromatic contrasts between deviant and standard stimuli for deuteranopia can be obtained. In deuteranopia, colors that align approximately along the red-green axis in typical trichromacy appear to be the same, forming confusion lines on a chromaticity diagram. According to Pridmore (2014), the empirical color axis in dichromats is orthogonal to the confusion line, intersecting the neutral gray point. As such, the chromatic distances between colors for deuteranopia can be represented as the distance along a line orthogonal to the confusion line passing through the neutral gray point D65 (Figure 1).

In addition, the confusion line intersecting D65 is considered as the categorical boundary for dichromacy (Broackes, 2010), where points on the same side of the boundary denote categorical similarity. Hence, the chromatic difference can be calculated as the sum of the orthogonal distances to the confusion line when the stimuli are positioned on opposite sides of the line. Conversely, when stimuli are on the same side, the chromatic difference is calculated by subtracting these distances.

The relative chromatic contrasts of the blue-green and red deviants to the green standard stimulus for deuteranopia were estimated to be 0.0290 and 0.0134, respectively. These estimations were based on the orthogonal distances of 0.0212, 0.0078, and 0.0078 for green, blue-green, and red, respectively, with the confusion line depicted in Figure 1. Therefore, blue-green was expected to be more salient than red when contrasted with green for individuals with deuteranopia and anomalous (deuteranomalous) trichromacy close to deuteranopia. However, this was not guaranteed for all anomalous trichromats, particularly for participants with a low red-green threshold (Table **??**). Categorically, red and green were expected to be more similar than blue-green and green in deuteranopia and anomalous trichromacy, with a high red-green threshold.

These theoretical estimates further predict a reversal in the saliency of deviant stimuli in typical trichromacy and deuteranopia because the ratio of blue-green to red to the distance from green was 0.03/0.052 = 0.57 for typical trichromacy, whereas it was 0.029/0.0134 = 2.16 for deuteranopia, although the absolute perceived contrasts were difficult to estimate.

Given the diversity of red-green thresholds across participants with anomalous trichromacy, chromatic contrasts between stimuli would naturally differ among participants. However, we prioritized the use of the same chromatic stimuli for all participants, rather than adjusting the chromatic contrast across participants. This approach allowed us to observe the neural diversity associated with differences in the perception of stimuli of the same color. To ensure that the luminance contrast was not used as a cue for the task, the luminance of the color stimuli was equated to 20 *cd/m*^2^ D65 for each participant using flicker photometry (See 2.5 Procedure for details).

### 2.3 Task Design

The oddball paradigm task (Sutton et al., 1965; Picton et al., 1992) was used as the attention-demanding task. This task comprised a frequently appearing standard stimulus and rarely appearing deviant stimulus presented in a random sequence. The participants were asked to focus on detecting the deviant stimulus (either blue-green or red) among the standard stimuli (green), and to respond by pressing a button immediately upon detecting the deviant stimulus (Figure 2). The choices of blue-green as deviant 1 and red as deviant 2 were based on the estimated chromatic contrasts from the standard stimulus, with the aim of invoking different levels of attention. Furthermore, the relationship of the chromatic contrast between these standard deviant pairs was designed to be reversed between typical trichromacy and deuteranopia (See 2.2 Stimuli).

**Figure 2.**
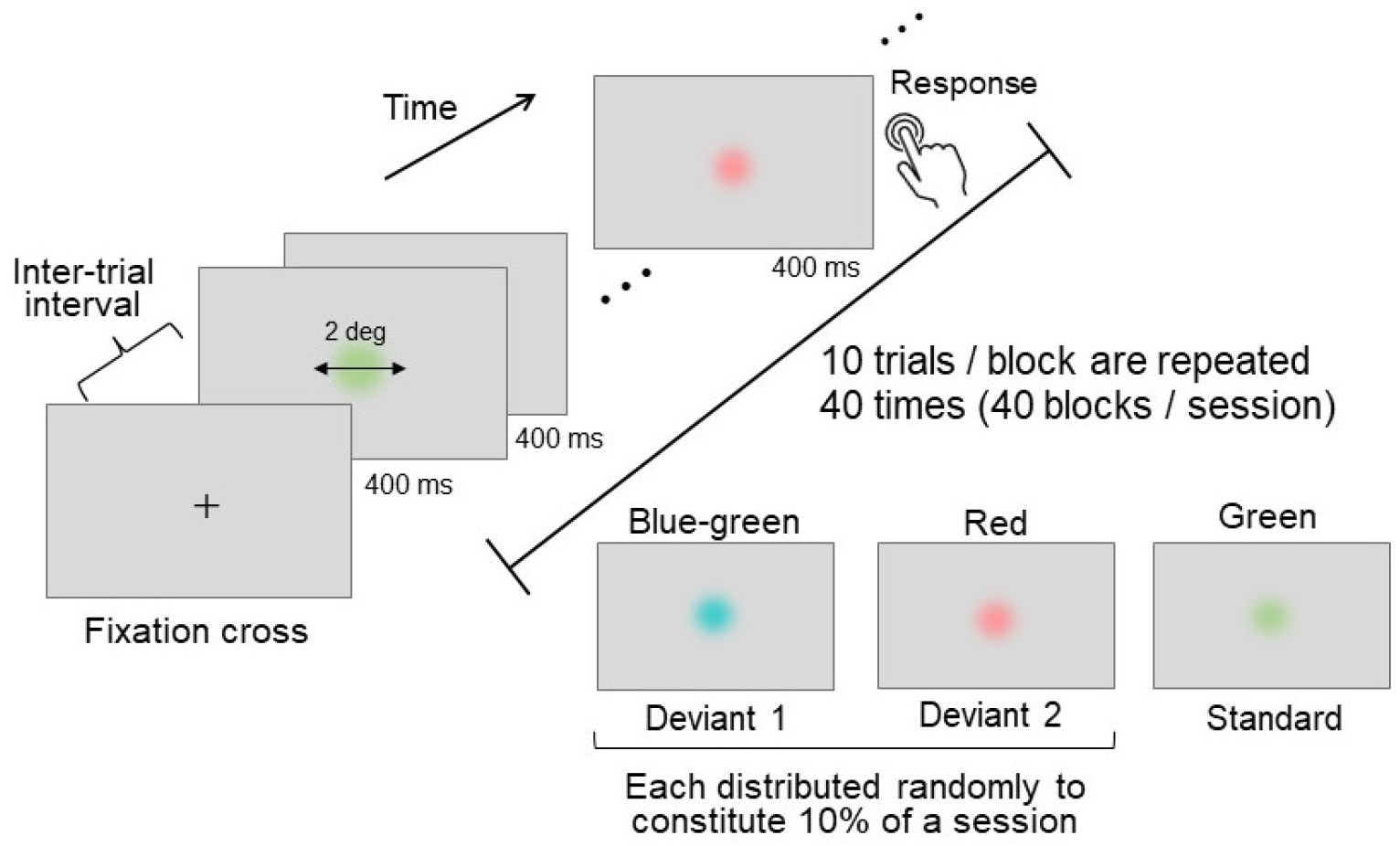
Schematics of the oddball task. Three distinct colors were used, each assigned either as a rarely presented deviant or high-frequent standard stimulus. Each stimulus was presented for a duration of 400 ms, with an inter-trial interval ranging between 1200–1600 ms. Participants were instructed to press a button immediately following detection of the deviant stimulus.

### 2.4 Apparatus

The stimuli were displayed on an LCD monitor (Display++, Cambridge Research Systems Ltd.), coupled with an image processor (Bits#, Cambridge Research Systems) capable of presenting colors at 14-bit resolution. The experimental program was controlled using MATLAB (MathWorks Inc.) and the psychtoolbox-3 extension (Brainard and Vision, 1997; Pelli and Vision, 1997; Kleiner et al., 2007). EEG signals were recorded using a 64-channel digital recorder (ActiCHamp Plus, Brain Products GmbH). Active electrodes were placed on the scalp following a 10–20 layout system utilizing an electrode cap (actiCap slim, Brain Products GmbH).

The timing of the stimulus onset and behavioral responses were recorded using an EEG amplifier equipped with a trigger box via a parallel port. The stimulus onset and stimulus color condition in each trial were detected using photoelectric sensors (MaP1180PS2A, NIHON SANTEKU Co.,Ltd) attached directly to small areas at the edge of the display and captured binary luminance modulations synchronized with the stimulus in these areas. The participants used a button connected to the trigger box to submit their behavioral responses.

### 2.5 Procedure

After placement of the electrode cap on their scalp, each participant was seated in a chair positioned 57 cm from the display surface with their eyes fixated at the center of the screen in a dark room. Prior to the oddball experiment, the luminance of the three stimulus colors (blue–green, red, and green) was adjusted to match 20 *cd/m*^2^ D65, the color used as a background, using flicker photometry (Wagner and Boynton, 1972; Kaiser and Comerford, 1975; Anstis and Cavanagh, 1983) with a 15 Hz alteration. Equiluminance measurements were conducted six times for each stimulus using the up-down method. The starting luminance was either 23 *cd/m*^2^ or 17 *cd/m*^2^, and the adjustment direction alternated between ascending and descending towards 20 *cd/m*^2^ D65 for each measurement. The resulting luminance values were averaged and used to set the stimulus luminance levels for subsequent oddball experiments.

In each oddball trial, a disk-shaped stimulus was presented at the center of the screen for 400 ms, with random inter-trial intervals ranging between 1200 and 1600 ms (Figure 2). The presented stimulus had a visual angle of 2^*°*^, and its edge was linearly and gradually blended into the gray D65 background to prevent the chromatic edge between the stimulus and background from influencing stimulus detection.

Each block consisted of ten consecutive trials. The 40 blocks were grouped into a single session and conducted twice for each participant. The presentation frequency ratio of the standard to deviant stimuli was 8:2, with an equal ratio between the two deviants. The trials featuring deviant stimuli were randomly distributed throughout each session. Although any given block could contain more than one deviant trial, because of randomization, not all blocks necessarily included deviant trials. To prevent confusion between stimuli, only one type of deviant stimulus, if included, was presented in a block. In addition, in each block, the appearance of a deviant stimulus was always preceded by a standard stimulus, ensuring that the discrimination of deviants was consistent with the standard stimulus. This systematic presentation facilitated the comparison of responses to deviant and standard stimuli.

To provide clear stimulus instructions and avoid uncertainty, color names were not used to designate stimulus types (standard or deviant). Instead, participants recognized deviant and standard stimuli by observing example stimulus sequences prior to the task. After identifying deviant and standard stimuli based on the appearance frequency difference, the participants completed three practice blocks to familiarize themselves with the task procedure. Following the practice phase, an experimental phase consisting of two sessions (80 blocks in total) was conducted.

During the experiment, the participants were allowed arbitrary interblock breaks, advancing to the next block by pressing the spacekey. Additionally, a mandatory 10-minute break between sessions was scheduled.

A total of 800 trials were conducted (640 standard stimulus trials [green], 80 deviant 1 [blue-green], and 80 deviant 2 [red] trials). During the trials, most participants used their right thumb to press the button to submit their reaction times (RTs) as soon as deviant stimuli were detected.

### 2.6 Behavioral Data analysis

The mean RTs for each deviant, including deuteranopia, were computed individually for each participant. Statistical differences between deviants were tested using paired *t*-tests, and effect sizes were calculated using Cohen’s *d* for both color vision groups (typical trichromats and anomalous trichromats). The difference between color vision types in each deviant condition was tested using a two-sample *t*-test. The significance level was set at *p <* 0.05. Multiple comparisons between the deviants and color vision types were corrected using the Bonferroni–Holm method (Holm, 1979). Owing to the limited sample size, individuals with deuteranopic vision were excluded from the group analysis, while the RTs between deviants were compared by paired *t*-tests based on the multi-trial data of the participant.

To examine the performance of the behavioral responses, hit rates were computed based on the average individual rate of responses to deviants 1 and 2. For the nontarget standard stimulus, false alarm rates were computed as the average of the individual false alarm rates.

Any RTs above 900 ms were identified as outliers and were excluded from the behavioral analysis. Statistical analyses were performed using MATLAB software (MathWorks, Inc.).

### 2.7 EEG Recording and Data Analysis

#### 2.7.1 Data Preprocessing

EEGs were recorded and digitized at a sampling rate of 1000 Hz. Subsequent analyses were conducted using EEGLAB (version 2022.0; Delorme and Makeig 2004) in MATLAB (MathWorks, Inc.). The data were organized following the Brain Imaging Data Structure (BIDS; Gorgolewski et al. 2016), with the extention of EEG data (Pernet et al., 2019). Initially, the data were bandpass-filtered between 0.5 Hz to 30 Hz. Two EEGLAB plugins, IC Label and Clean Rawdata, were employed to enhance data readability and quality. The IC Label function differentiates independent components (ICs) stemming from brain activity from non-brain sources. ICs related to eye movements, muscle artifacts, or specific channels with greater than 90 % probability were marked for rejection. In addition, the Clean Rawdata plugin identified and removed data segments and channels that were significantly contaminated by noise. We adhered to the rejection criteria recommended by Delorme (2023), and supplemented this process by interpolating deleted channels with data from neighboring preserved channels, thereby optimizing the data quality before referring to the average. In the final preprocessing step, 1000 ms of individual trial data were extracted from the recordings. The average of the 100 ms pre-stimulus period was subtracted to establish the 0 *µ*V baseline (Murray et al., 2008; Luck, 2014; Keil et al., 2014).

#### 2.7.2 Data Analysis

The EEG data were segmented into single-trial data with a duration of 1000 ms post-stimulus onset. These segments were averaged for individual participants and further grand-averaged across participants to compute the ERPs for each stimulus condition within each color vision type (typical and anomalous trichromats). In addition, 95 % confidence intervals (CIs) were computed to present the variation among participants. To assess stimulus-induced brain activity, the average ERPs at the centroparietal electrodes (Cz, CPz, and Pz) were visually inspected. Given that the visual oddball task is an attention-demanding task, typical attention-related ERP components, such as P3, were expected to be observed from these electrodes (Zhang and Kappenman, 2024), if the experimental paradigm worked successfully. P3 is a late positive component that typically appears approximately 400 ms after post-stimulus onset in response to rare stimulus condition within a sequence of stimuli. This tends to be heightened toward more difficult stimuli (Polich, 1987; Alho et al., 1992), reflecting an increase in the allocation of attentional resources (Grasso et al., 2009; Isreal et al., 1980). Average ERPs at the frontal (AF3, AFz, and AF4) and occipital (PO7, O1, O2, and PO8) electrodes were visually inspected to confirm the time course at these scalp locations.

During the oddball task, differences in the perceptual and cognitive processes related to color saliency and color vision types may manifest as distinct spatiotemporal patterns in neural activity. To comprehensively explore these patterns, we employed an exploratory statistical analysis technique known as cluster-based permutation analysis. This approach is nonparametric and data-driven, allowing us to assess the differences in neural activity across multiple time points and channels without predefined assumptions regarding specific time or spatial locations with nominal Type I error rates (Maris and Oostenveld, 2007; Sassenhagen and Draschkow, 2019).

The cluster-based permutation test evaluates the null hypothesis that different conditions (e.g., different stimulus conditions or color vision types) are sampled from the same distribution; therefore, they are inter-exchangeable. If the observed effect was unlikely (less than 2.5 % in the two-tailed test) under label-shuffled data, the hypothesis was rejected, indicating that the observation of the effect was not by chance. Using this method, we compared i) the stimulus color conditions for each color vision type and ii) the color vision types under the same stimulus color conditions. We utilized a set of statistical functions provided by Fieldtrip (Oostenveld et al., 2011) to perform a cluster-based permutation test. Individual mean ERPs across the 64 electrodes during the 600 ms following stimulus onset were used for analyses. This time window was selected to analyze both perceptual and cognitive neural activity, covering the P3 ERP component, a late-positive ERP component related to attentional processes.

A cluster is defined as the sum of *t*-statistics grouped together based on temporal and spatial adjacency. In our analysis, a cluster was formed with at least two continuous time points and/or neighboring electrodes that exhibited statistical significance above the critical alpha level of 0.05. The cluster threshold was calculated based on a cluster distribution of 10,000 random partitions using the Monte Carlo method. The cluster size (sum of *t*-values) with occurrences of probabilities of less than 0.625 % (2.5 % with Bonferroni correction for the four repeated comparisons) on either side of the tail was identified to determine the significance threshold. The significance of the detected clusters was evaluated on the basis of the determined thresholds. Clusters with positive *t*-values were considered positive clusters, whereas clusters with negative *t*-values were considered negative clusters. Cohen’s effect size was computed based on the mean potential changes during the period of the observed significant clusters.

### 3 RESULT

#### 3.1 Flicker Photometry

The luminance of the stimuli was individually adjusted to be equiluminant with a gray background before the oddball task using flicker photometry. The mean luminance values for deviant 1, deviant 2, and standard stimuli were 20.49 *±* 0.36, 19.74 *±* 0.37, and 20.11 *±* 0.12 *cd/m*^2^ for typical trichromats, 21.27 *±* 0.086, 18.96 *±* 0.12, and 20.60 *±* 0.095 *cd/m*^2^ for anomalous trichromats, and 21.43 *±* 0.30, 19.04 *±* 0.32, and 20.77 *±* 0.58 *cd/m*^2^ for the deuteranope, respectively.

All three stimuli, deviant 1 (*p* = 0.0003, Cohen’s *d* = 2.35), deviant 2 (*p* = 0.0003, Cohen’s *d* = 2.3), and standard (*p* = 0.024, Cohen’s *d* = 1.25), presented statistically significant differences between color vision types (typical trichromats and anomalous trichromats).

### 3.2 Behavioral Results

In Figure 3, individual mean RTs are presented in two separate plots: one for typical trichromats and another for the anomalous trichromats and deuteranope. For typical trichromats, the mean RT for deviant 2 was significantly shorter than that for deviant 1 (deviant 1: 393 *±* 78 ms; deviant 2: 382 *±* 78 ms; *p*_*adj*_ = 0.002, Cohen’s *d* = 0.14). However, there was no significant difference between deviants 1 (414 *±* 82 ms) and 2 (418 *±* 86 ms) for anomalous trichromats (*p*_*adj*_ = 0.39, Cohen’s *d* = 0.06). Typical trichromats had significantly faster RTs for both deviants than anomalous trichromats (deviant 1: *p*_*adj*_ *<* 0.0001, Cohen’s *d* = 0.27; deviant 2: *p*_*adj*_ *<* 0.0001, Cohen’s *d* = 0.45).

**Figure 3.**
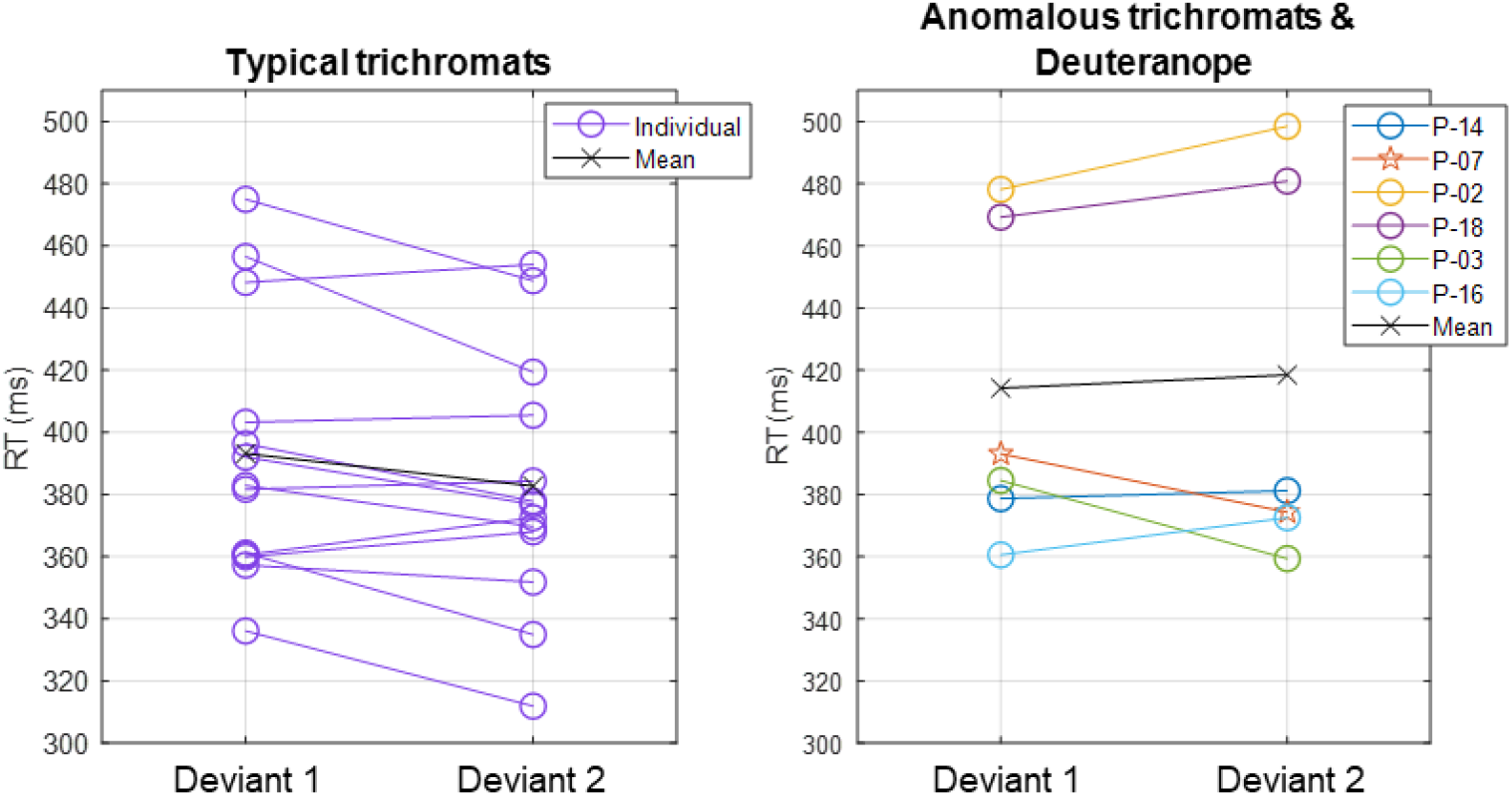
Individual mean RTs for each participant and group mean RTs for typical trichromats and anomalous trichromats for deviants 1 and 2. For anomalous trichromats, the colors of the symbols correspond to the participants outlined in Table 1, in which participants are listed in descending order according to their red-green threshold, as are listed in the legend of the right panel. Stars, plotted with anomalous trichromats, indicate individual mean RTs for the deuteranope while circles indicate those for typical and anomalous trichromats.

Participants with deuteranopia exhibited a significantly faster RT to the deviant 2 (374 *±* 57 ms) than deviant 1 (393 *±* 58 ms), demonstrating a faster response to the less salient deviant (*p* = 0.04, Cohen’s *d* = 0.33).

**Table 1.**
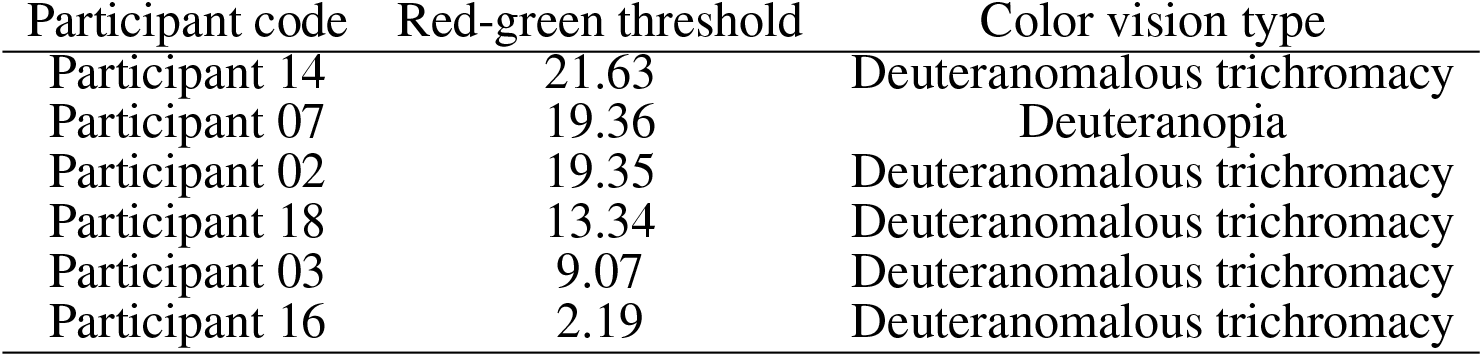
Red-green thresholds as assessed using the CAD test, and color vision types identified based on anomaloscope test for the participants with minority color vision phenotypes. Participants are listed in descending order according to their red-green threshold.

| Participant code | Red-green threshold | Color vision type |
| --- | --- | --- |
| Participant 14 | 21.63 | Deuteranomalous trichromacy |
| Participant 07 | 19.36 | Deuteranopia |
| Participant 02 | 19.35 | Deuteranomalous trichromacy |
| Participant 18 | 13.34 | Deuteranomalous trichromacy |
| Participant 03 | 9.07 | Deuteranomalous trichromacy |
| Participant 16 | 2.19 | Deuteranomalous trichromacy |

Hit rates, which reflect the rates of successful responses to deviants, were high across all color vision types. The group mean hit rates were 99.71 % and 99.42 % for typical trichromats, 100 % and 99.50 % for anomalous trichromats, and 100 % for one deuteranope participant for deviants 1 and 2, respectively. The group mean false alarm rates, which provided the likelihood of responses to the non-target standard stimulus, were low for all color vision types: 0.1 % for typical trichromats, 0.05 % for anomalous trichromats, and 0.17 % for deuteranopes. Due to few recorded false trials, it was impossible to identify which deviant was more likely to be falsely detected.

Three RT records were identified as outliers for typical trichromats: one from deviant 1 (919 ms) and two from deviant 2 (1588 and 1069 ms). All the three records included the same participants. Similarly, three RT records were identified as outliers from anomalous trichromats; once from deviant 1 (986 ms) and twice from deviant 2 (1051 and 981 ms); again the records belonged to the same participant. These records were therefore excluded from the behavioral analyses.

### 3.3 Electrophysiological Result

#### 3.3.1 ERPs

Figure 4 shows the ERPs of each color vision group in the three distinct scalp regions. The central column displays ERPs in the parietal region, averaged across Cz, CPz, and Pz. Notably, the ERP component P3 was observed in both deviant conditions, whereas the response to the standard condition was suppressed in both color vision groups. The 95 % confidence interval for anomalous trichromats tended to be wider than that for typical trichromats.

**Figure 4.**
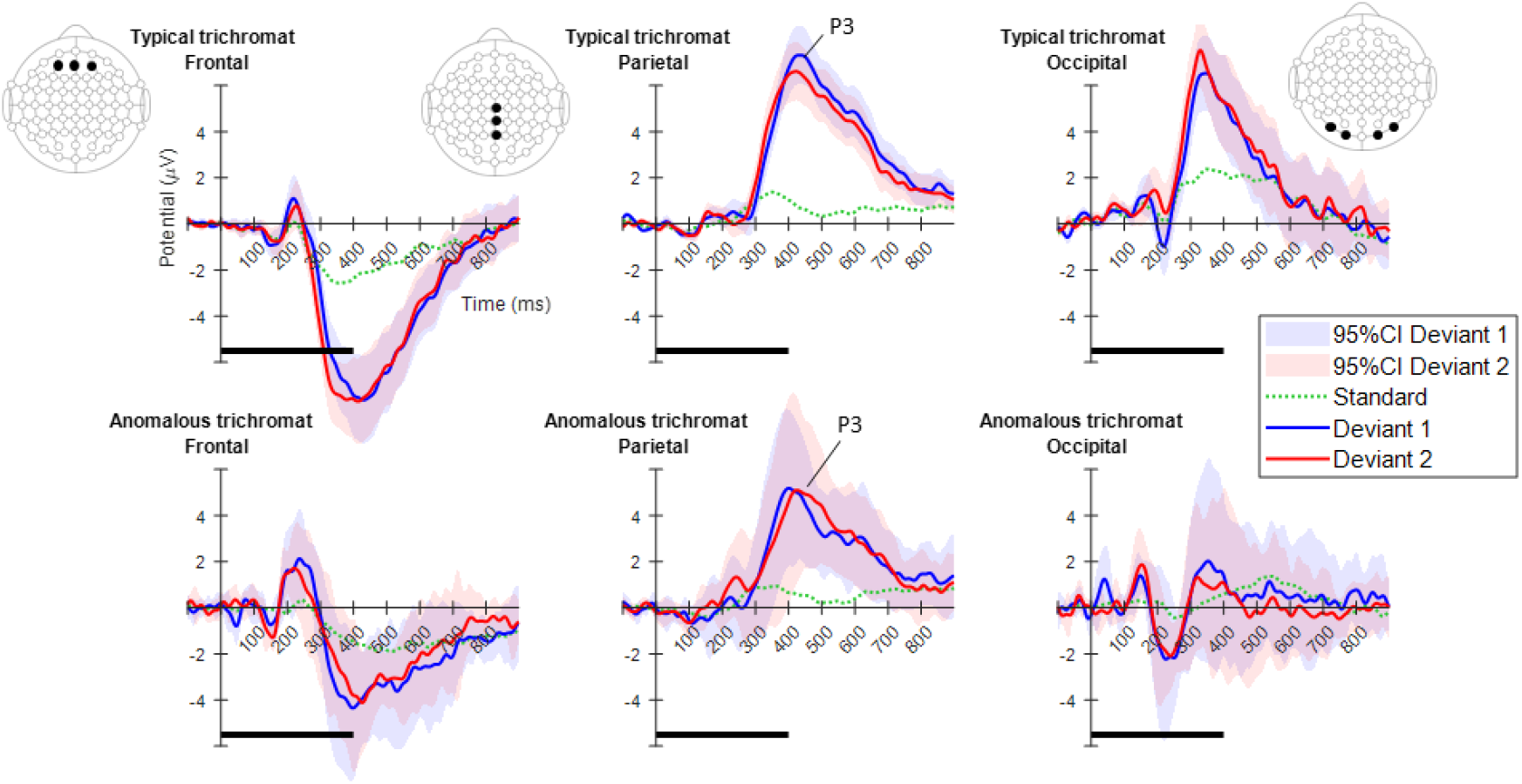
Mean ERPs for each color vision group in the three scalp regions. From left to right, ERPs are shown for the frontal, parietal, and occipital regions. The electrode positions included in each region are highlighted using black dots in the electrode layout. The plots in the top row indicate typical trichromats, while those in the bottom row indicate anomalous trichromats. In each plot, blue and red lines represent the mean ERPs across participants for deviants 1 and 2, while shaded areas in the same color indicate 95 % confidence intervals (CIs) for each deviant. A dotted green line indicates the mean potential for standard stimulus. Horizontal black lines at the bottom of each plot indicate the duration of stimulus presentation.

#### 3.3.2 Exploratory analysis of ERPs

Cluster-based permutation tests show a significant difference between the deviant conditions with a negative cluster for typical trichromats (positive cluster: *p*_*adj*_ = 0.035, Cohen’s *d* = 0.70; negative cluster: *p*_*adj*_ = 0.003, Cohen’s *d* = 0.58). The cluster indicating the higher amplitude for deviant 2 in the analyzed time window extended from approximately 200 to 350 ms post stimulus and was predominantly distributed in the occipital region up to 240 ms, before gradually spreading and shifting toward the parietal region, exhibiting a maximal response difference at approximately 300 ms (Figure 5a).

**Figure 5.**
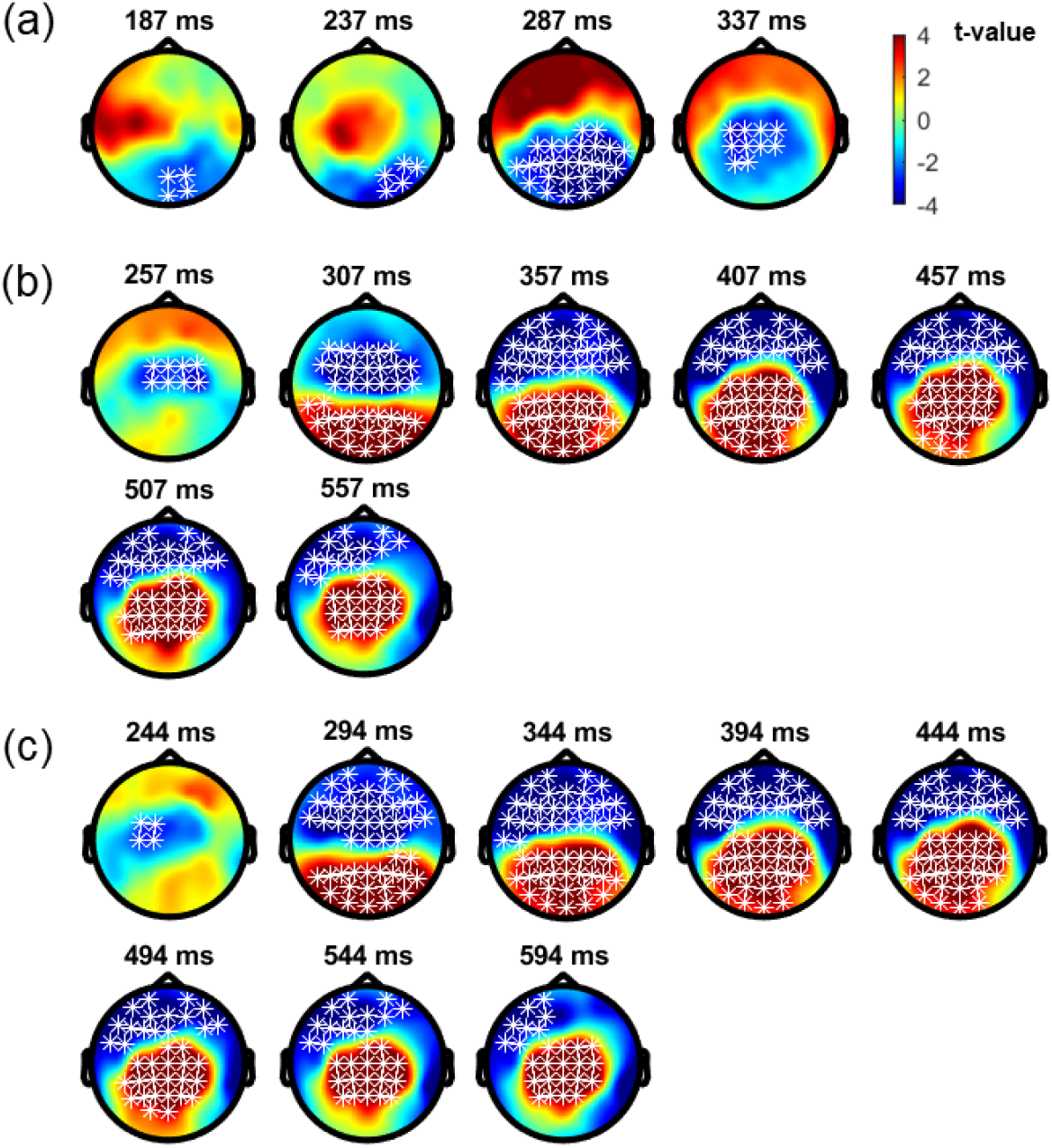
Results of the cluster-based permutation analysis for typical trichromats. (a) Comparison between deviant 1 and deviant 2 conditions. (b) Comparison between deviant 1 and standard conditions. (c) Comparison between deviant 2 and standard conditions. The distribution of textitt-values are topographically plotted, with electrodes highlighted with white asterisks to indicate locations of the clusters with 50 ms interval.

Comparisons of each deviant condition with the standard condition also revealed significant differences in typical trichromats (Figure 5b, c). When comparing deviant 1 with the standard, a positive cluster indicating a higher amplitude for deviant 1 appeared around the occipital and parietal regions (*p*_*adj*_ = 0.0004, Cohen’s *d* = 2.9), and a negative cluster indicating a lower amplitude for deviant 1 appeared around the frontal region (*p*_*adj*_ = 0.0008, Cohen’s *d* = 1.7). Similar clusters were observed in the comparison of deviant 2 with the standard (positive cluster: *p*_*adj*_ = 0.0004, Cohen’s *d* = 2.2; negative cluster: *p*_*adj*_ = 0.004, Cohen’s *d* = 1.5). These clusters appeared approximately 250 ms post stimulus and lasted until the end of the analysis window for both comparisons.

There were no significant differences in anomalous trichromats between the deviant conditions (positive cluster: *p*_*adj*_ = 0.12, Cohen’s *d* = 0.64; negative cluster: *p*_*adj*_ = 0.88, Cohen’s *d* = 1.0). A significant difference was found when comparing deviant 1 with the standard. A positive cluster indicating a higher amplitude for deviant 1 appeared around the occipital to parietal regions approximately after 350 ms post stimulus, lasting until the end of the analysis window (positive cluster: *p*_*adj*_ = 0.0004, Cohen’s *d* = 3.3; negative cluster: *p*_*adj*_ = 0.13, Cohen’s *d* = 3.1) (Figure 6a). Similarly, a significant difference was observed when comparing deviant 2 with the standard, with a positive clusters indicating a higher amplitude for deviant 2 appeared around the occipital to parietal regions approximately 400–500 ms post stimulus (positive cluster: *p*_*adj*_ = 0.0004, Cohen’s *d* = 3.4; negative cluster: *p*_*adj*_ = 0.12, Cohen’s *d* = 3.8).

**Figure 6.**
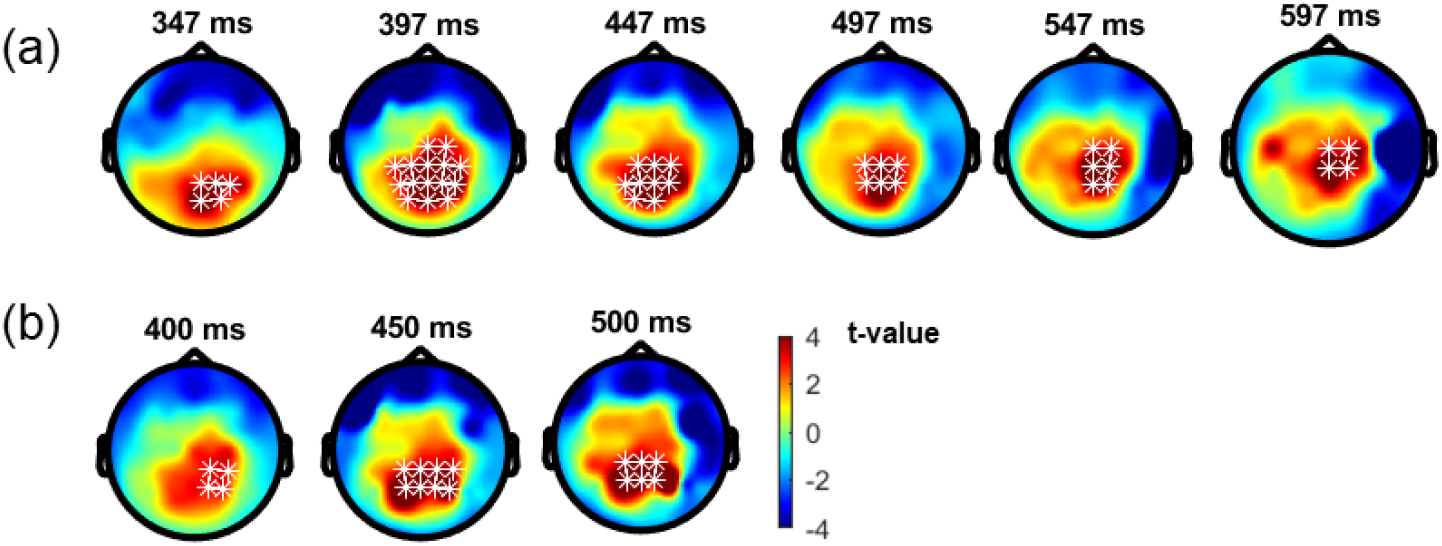
Results of the cluster-based permutation test for anomalous trichromats. (a) Comparison between deviant 1 and standard conditions. (b) Comparison between deviant 2 and standard conditions. The distribution of textitt-values are topographically plotted, with electrodes highlighted with white asterisks to indicate locations of the clusters with 50 ms interval.

When comparing the data of typical and anomalous trichromats using a cluster-based permutation test, no significant differences were found for either the deviant condition (deviant 1 positive cluster: *p*_*adj*_ = 1, Cohen’s *d* = 1.9; deviant 1 negative cluster: *p*_*adj*_ = 1, Cohen’s *d* = 1.9; deviant 2 positive cluster: *p*_*adj*_ = 0.14, Cohen’s *d* = 2.0; deviant 2 negative cluster: *p*_*adj*_ = 0.66, Cohen’s *d* = 2.0).

## 4 DISCUSSION

In this study, we investigated the neural activity in response to variations in color saliency across different color vision types during an attention-demanding oddball task. The aim of this investigation was to elucidate the spatiotemporal characteristics of the neural activity underlying the perceptual and cognitive processes involved in this task among different color vision types, with a particular focus on anomalous trichromats. Specifically, we sought to explore the neural activities that may bridge the gap between red-green sensitivity and cognitive behavior.

To facilitate the examination of commonalities and differences in responses under uniform conditions, consistent chromaticity of color stimuli was used across participants. During the oddball task, two deviant stimuli (blue-green and red) were employed to investigate the reverse saliency conditions between typical trichromats and most anomalous trichromatic participants.

We expected typical trichromats to demonstrate faster RTs to the more salient red stimulus compared to the less salient blue-green stimulus, along with the neural activity reflecting increased attentional demand for the less-salient blue-green stimulus. Additionally, due to the variability in red-green perceptual sensitivities among anomalous trichromats, both neural and behavioral responses were expected to reflect sensitivity diversity under the reversed-stimulus condition compared to typical trichromats.

Observation of the P3 component over the parietal region for both deviant stimuli confirmed that participants directed their attention toward these deviant stimuli, while discerning them from the standard stimulus, as instructed. Exploratory ERP analysis revealed a cluster indicating higher potentiation of the red stimulus than the blue-green stimulus around the occipital to parietal regions, accompanied by lower potentiation of the red stimulus around the frontal region. Although this pattern initially appeared inconsistent with our expectations, cluster-based permutation analysis revealed a consistent temporal trend of faster neural amplification for the more salient red stimulus during the P3 response, validating our experimental paradigm.

In contrast to typical trichromats, the effect of saliency differences was ambiguous in anomalous trichromats, showing a larger variability in both behavioral and neural responses. For instance, a participant with deuteranopic dichromacy exhibited faster RTs to a red stimulus, which was expected to be less salient to someone with a higher red-green threshold. Despite generally longer RTs compared to typical trichromats, both dichromats and anomalous trichromats maintained high hit rates and low false alarm rates for both deviant stimuli, indicating successful stimulus identification. These results indicate that perceptual and cognitive factors, including individual strategic differences, influence RTs in participants with minority color vision phenotypes. Nevertheless, our results are consistent with previous findings showing a weak correlation between cone spectral sensitivity and perceptual chromatic sensitivity in anomalous trichromats (Bosten, 2019). The observed RTs underscore the complexity and variability of color perception among individuals with minority color vision phenotypes.

The neural processes underlying these phenomena may offer insights into the intricate factors bridging the gaps between receptoral, perceptual, cognitive, and behavioral aspects in color vision research. While our exploratory analysis of ERPs did not unveil any significant differences between deviant conditions in anomalous trichromats, an examination of ERPs elicited by the green standard stimulus revealed temporal differences in neural activity between conditions with differing saliency. During the deviant-standard comparison analysis, the cluster reflecting higher potentiation to blue-green appeared at approximately 350 ms post stimulus, whereas the cluster reflecting higher potentiation to red appeared at approximately 400 ms post stimulus, predominantly in the occipital region and extending to the parietal region. Although the results of cluster-based permutation tests do not precisely pinpoint the timings and locations of distinct neural activities between conditions (Sassenhagen and Draschkow, 2019), this slight discrepancy in timing between deviant conditions may be linked to previously reported faster RTs in visual search tasks among individuals with anomalous trichromacy (Sunaga et al., 2013), in which faster RTs to blue-green were reported amidst green and blue distractors.

It should be noted that large individual differences in ERP waveforms were observed in both typical and anomalous trichromats (see Supplementary Materials). However, due to individual variations in physiological factors such as skull thickness, the extent to which these variations reflect differences in neural activity remains unclear (Hakim et al., 2021). Additionally, it is important to acknowledge that EEG signals primarily capture the activity of pyramidal neurons, with minimum contribution from interneuron activity (Luck, 2014). The observed individual variability in neural activities, along with inherent technical limitations of EEG measurements and differences in experimental paradigms, may have contributed to the challenges in observing neural activity reflecting the enhanced sensitivity in the early visual cortex in anomalous trichromats, as reported in an fMRI study (Tregillus et al., 2021). Moreover, cluster-based permutation tests tend to be insensitive to localized activities. In this study, relatively smaller deflections were observed during an earlier time window prior to P3, which was more localized than the P3 activity. Notably, potential conditional differences during this earlier time window may have been overlooked in the analysis.

Another limitation of this study lies in the uncertainty regarding the origin of the variation in red-green sensitivity and whether it arises from inherent genetic factors or neuroplastic changes developed through ontogeny. Clarifying the relationship between perceptual diversity and variations in neural activity requires the untangling of genetic factors, such as those related to cone sensitivities and developmental influences, or exploring their interplay. Integrating these factors into future studies will be critical to obtain a more comprehensive understanding of the relationship between perceptual and neural diversity.

## 5 CONCLUSION

In the present study, we investigated the spatiotemporal dynamics of neural activity corresponding to chromatic differences in both anomalous and typical trichromats. The results revealed discernible differences in neural activity between typical trichromats. Anomalous trichromats demonstrated a similar temporal pattern, showing faster neural responses toward a color that was expected to be more salient under color reversal conditions. However, no obvious differences were found in the neural responses between the two groups. While both groups exhibited low behavioral error rates, only typical trichromats demonstrated consistent alignment between behavioral and neural responses. These findings highlight the complex relationship between saliency differences and neural representations of anomalous trichromacy. Further studies with larger sample sizes are required to better characterize the neural activity associated with the perceptual, cognitive, and behavioral processing of various color vision types.

## CONFLICT OF INTEREST STATEMENT

The authors declare that this study was conducted in the absence of any commercial or financial relationships that could be construed as a potential conflict of interest.

## AUTHOR CONTRIBUTIONS

NT: methodology, data collection, formal analysis, funding acquisition, and writing— original draft preparation. XC: data collection. HT, SM, MS, and YM: Methodology. CH: conceptualization, methodology, project administration and supervision, and funding acquisition. All authors contributed to the review and editing and have read and agreed to the published version of the manuscript.

## FUNDING

This work was supported by JSPS KAKENHI Grant Numbers JP19H04198 and JP23K17643, and the NTT-Kyushu University Collaborative Research Program for Fundamental Sciences (to C.H.). Support was further provided by JST SPRING, Grant Number JPMJSP2136 (to N. T.).

## ACKNOWLEDGMENTS

We would like to thank Daisuke Koike and Hiroshi Hashimoto for their contributions to the experiments. We are grateful to Shin’ya Nishida, Takahiro Kawabe, and Satoshi Nakadomari for constructive comments and general discussion on this study.

## Notes

### Competing Interest Statement

The authors have declared no competing interest.

### Summary of Updates

All sections of the previous version were revised based on new analyses. Specifically, utilizing multi-channel measured EEG, an exploratory analysis of neural data using cluster-based permutation tests was conducted to examine differences in spatiotemporal activity patterns by color vision types and color stimuli. This revision alleviates the concerns of the previous results due to the small sample size of participants with minority color vision. The new analysis provided statistically robust results and improved discussion.

