## Supplementary Material for "Temporal and Spatial Analysis of Event-Related Potentials in Response to Color Saliency Differences Among Various Color Vision Types"

### 1 SUPPLEMENTARY FIGURES

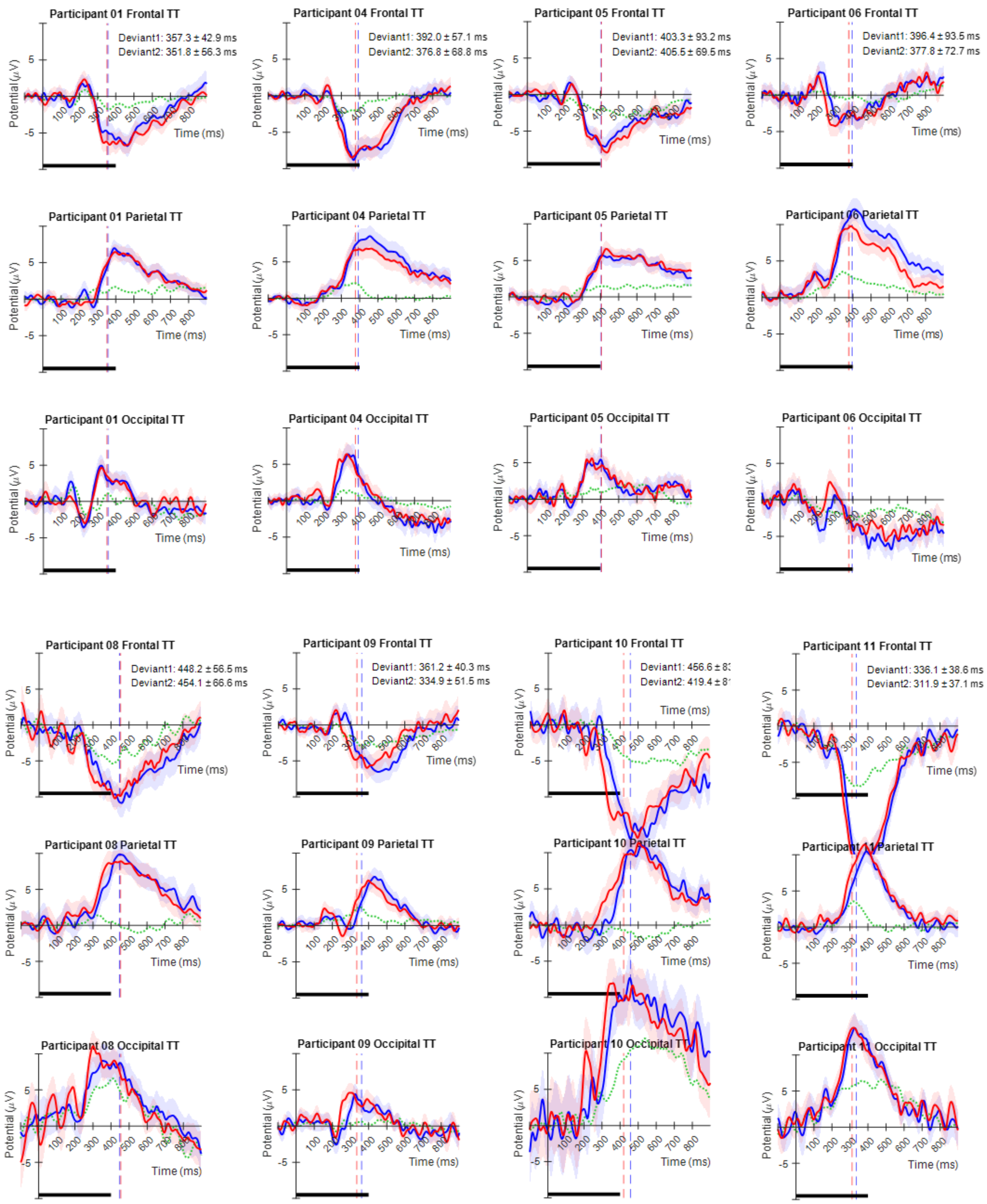
Participant 01 Frontal TT
Deviant1: 357.3 ± 42.9 ms
Deviant2: 351.8 ± 56.3 ms
Participant 04 Frontal TT
Deviant1: 392.0 ± 57.1 ms
Deviant2: 376.8 ± 68.8 ms
Participant 05 Frontal TT
Deviant1: 403.3 ± 93.2 ms
Deviant2: 405.5 ± 69.5 ms
Participant 06 Frontal TT
Deviant1: 396.4 ± 93.5 ms
Deviant2: 377.8 ± 72.7 ms
Participant 01 Parietal TT
Participant 04 Parietal TT
Participant 05 Parietal TT
Participant 06 Parietal TT
Participant 01 Occipital TT
Participant 04 Occipital TT
Participant 05 Occipital TT
Participant 06 Occipital TT
Participant 08 Frontal TT
Deviant1: 448.2 ± 56.5 ms
Deviant2: 454.1 ± 66.6 ms
Participant 09 Frontal TT
Deviant1: 361.2 ± 40.3 ms
Deviant2: 334.9 ± 51.5 ms
Participant 10 Frontal TT
Participant 11 Frontal TT
Deviant1: 336.1 ± 38.6 ms
Deviant2: 311.9 ± 37.1 ms
Participant 08 Parietal TT
Participant 09 Parietal TT
Participant 10 Parietal TT
Participant 11 Parietal TT
Participant 08 Occipital TT
Participant 09 Occipital TT
Participant 10 Occipital TT
Participant 11 Occipital TT
Potential (μV)
Time (ms)

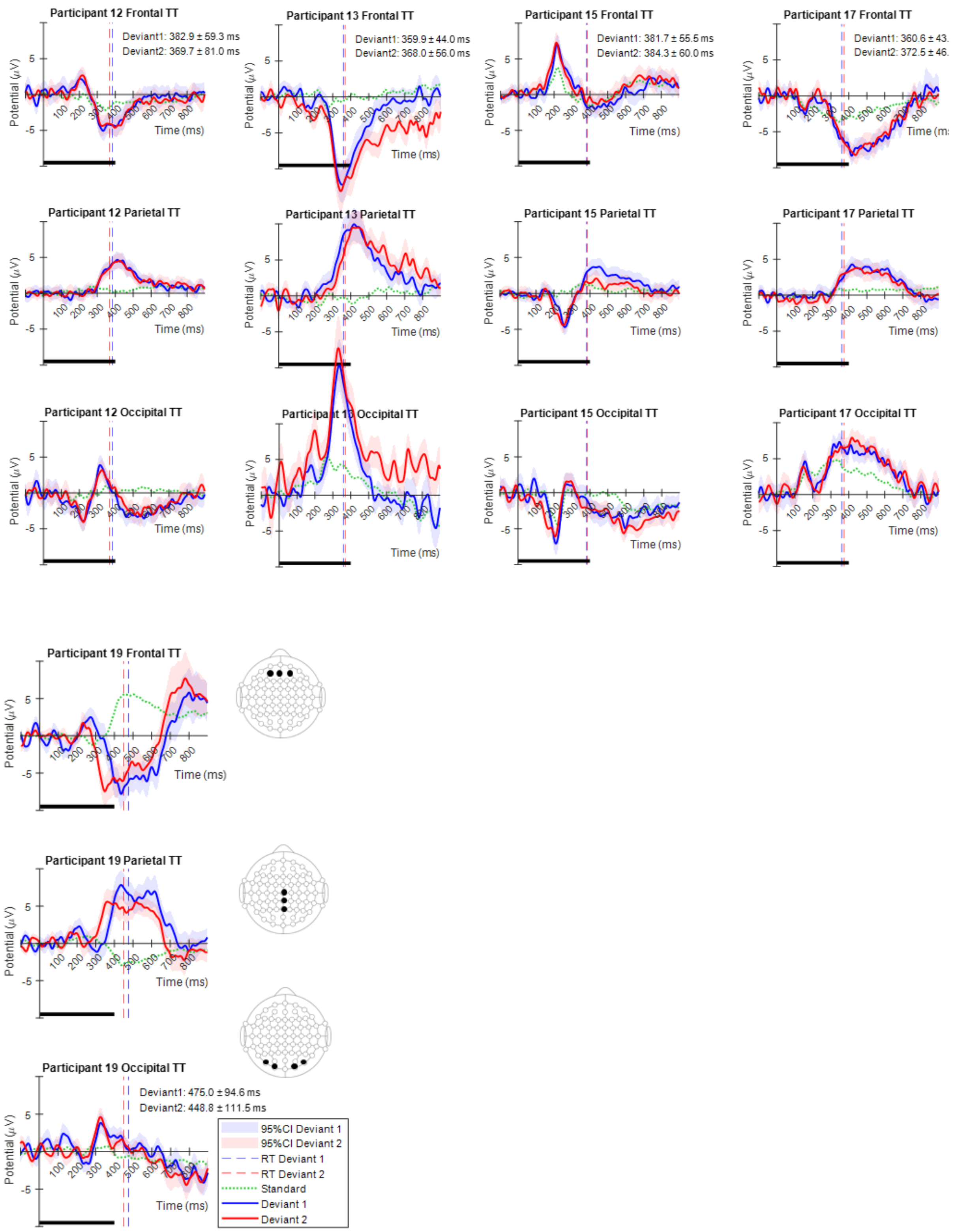


Figure S1: Individual ERPs from typical trichromats (TT). Post-preprocessed EEG data from the frontal, parietal, and occipital regions were averaged over trials for each individual. Averaged electrodes for each region correspond to AF3, AFz, AF4 in the frontal region, Cz, CPz, Pz in the parietal region, and PO7, O1, O2, PO8 in the occipital region. The mean and standard deviation of the RTs for each deviant stimulus are indicated as text inside the plot. The thick black line at the bottom indicates the stimuli presentation period.

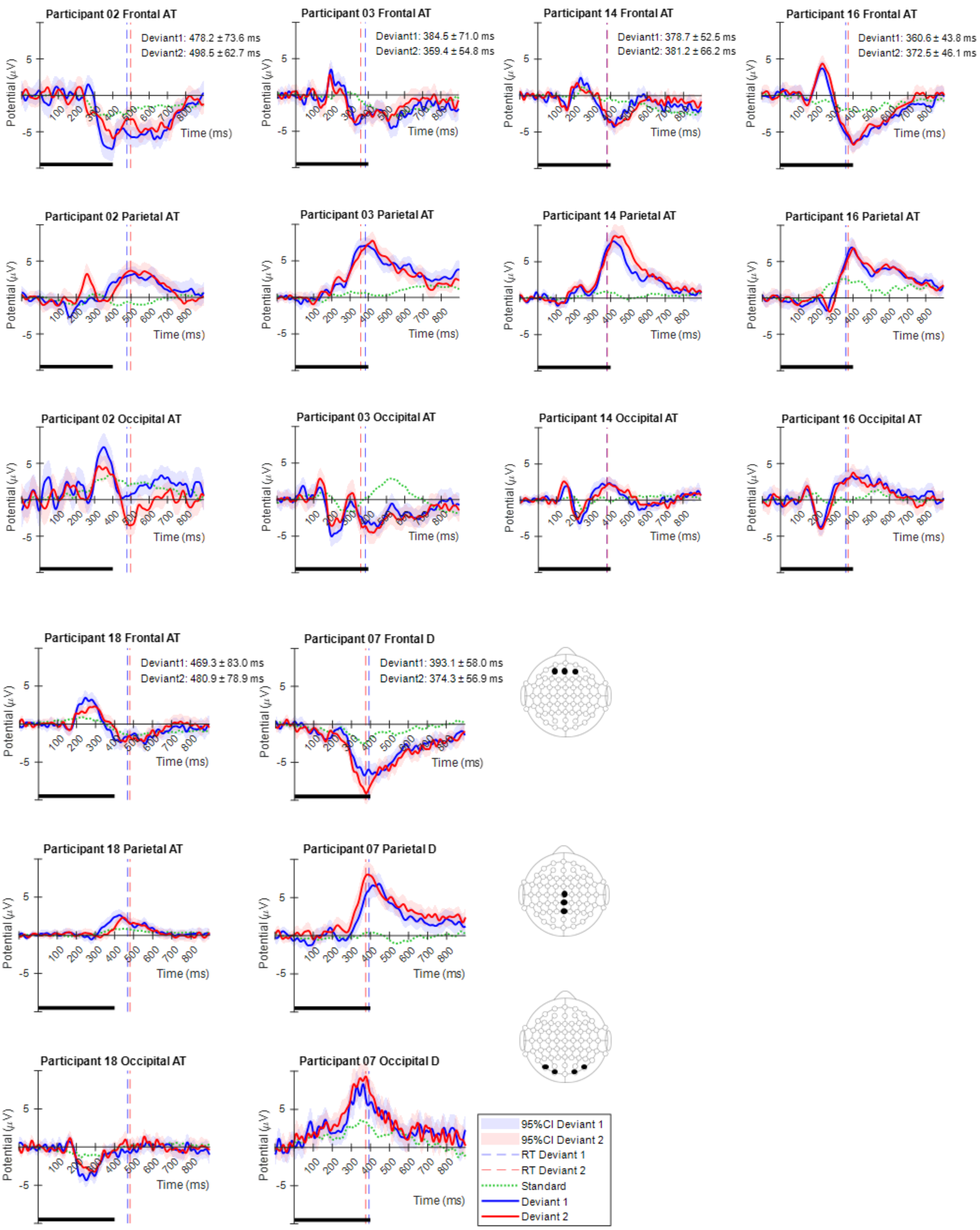


Figure S2: Individual ERPs from anomalous trichromats (AT) and deuteranopic dichromats (D). Post-preprocessed EEG data from the frontal, parietal, and occipital regions were averaged over trials for each individual. The averaged electrodes for each region correspond to AF3, AFz, AF4 in the frontal region, Cz, CPz, Pz in the parietal region, and PO7, O1, O2, PO8 in the occipital region. The mean and standard deviation of the RTs for each deviant stimulus are indicated as text inside the plot. The thick black line at the bottom indicates stimuli presentation period. Participant number 07 had deuteranopic dichromacy, while the rest had anomalous trichromacy (all are deuteranomalous trichromacy).
